# Social selection parapatry in an Afrotropical sunbird

**DOI:** 10.1101/027417

**Authors:** Jay P. McEntee, Joshua V. Peñalba, Chacha Werema, Elia Mulungu, Maneno Mbilinyi, David Moyer, Louis Hansen, Jon Fjeldså, Rauri CK Bowie

## Abstract

The extent of range overlap of incipient and recent species depends on the type and magnitude of phenotypic divergence that separates them. Trait divergence by social selection likely initiates many speciation events, but may yield niche-conserved lineages predisposed to limit each others’ ranges via ecological competition. Here we examine this neglected aspect of social selection speciation theory in relation to the discovery of a non-ecotonal species border between sunbirds. We find that *Nectarinia moreaui* and *N. fuelleborni* meet in a ~6 km wide contact zone, as estimated by molecular cline analysis. These species exploit similar bioclimatic niches, but sing highly divergent learned songs, consistent with divergence by social selection. Cline analyses suggest that within-species stabilizing social selection on song-learning predispositions maintains species differences in song despite both hybridization and cultural transmission in the contact zone. We conclude that ecological competition between *moreaui* and *fuelleborni* contributes to the stabilization of the species border, but that ecological competition acts in conjunction with reproductive interference. The evolutionary maintenance of learned song differences in a hybrid zone recommend this study system for future studies on the mechanisms of learned song divergence and its role in speciation.

## Background

The types of divergence processes that separate incipient or recent species should determine their abilities to limit each others’ ranges [1]. Whereas niche divergence can permit co-existence by easing ecological competition, divergence under niche conservatism may predispose incipient species to mutual range limits where contact occurs. Mutual range limits (hereafter parapatry) should therefore develop between incipient species divergent primarily by genetic incompatibility from mutation-order differences, reproductive trait divergence via random genetic drift, and/or social selection (inclusive of sexual selection). Parapatric boundaries involving minimally ecological divergence should form even where ecological gradients are shallow or absent, as their existence does not depend on advantages accrued via local adaptation on either side of the boundary [2-4]. Additionally, because biological factors trump abiotic factors in limiting ranges more frequently at lower latitudes in terrestrial environments [5], parapatric boundaries following speciation under niche conservatism should occur disproportionately in the tropics.

West-Eberhard [6] hypothesized that social selection-driven speciation in particular should frequently contribute to speciation events and predispose its products to parapatric distributions. According to this “social selection parapatry” hypothesis, the lag in ecological divergence during social selection-driven speciation should prevent co-existence of forms because strong resource competition occurs when incipient or recent species interact. Taxa subject to social selection divergence should be especially prone to both interspecific interference competition and preferential aggregation with conspecifics [6-9], and that these phenomena should also stabilize parapatry. While the roles of both social selection and niche conservatism [10,11] in speciation have recently been central foci of research, the particulars of West-Eberhard’s social selection parapatry hypothesis have received little attention. The absence of proposed examples may be due to the large number of component sub-hypotheses and corresponding required evidence set, or because detailed studies of non-ecotonal parapatric distributions from the tropics, where such distributions should be frequently encountered, are uncommon (but see [12]).

Here we examine the divergence and shared range boundary between the sunbird sibling species *Nectarinia moreaui* and *N. fuelleborni* in southern Tanzania in the context of the social selection parapatry hypothesis. These taxa exhibit similar habitat requirements, being found exclusively in “sky island” montane forests in eastern Africa. Unusually, their parapatric boundary occurs within one such sky island, the Udzungwa Mountains. Though only subtle differences in plumage exist [13], we find that males of the two species sing strikingly different territorial songs. Their abutting distributions present the opportunity to examine the processes that stabilize the species border of social trait-diverged sibling species with similar habitat requirements.

The meeting of recent or incipient species with highly divergent learned song moreover provides the opportunity to investigate learned song divergence’s role in speciation. Learned song has received a great deal of attention as a social trait that may be important to bird speciation [14-18], both because song-learning birds are especially diverse and because learned songs appear especially labile. Essential for song to have a central role in speciation, though, is its robustness to convergence during secondary contact of divergent speciesn [19-22]. If learned song differences help prevent the merging of incipient species, accrued differences must be maintained when interspecific interactions make possible heterospecific cultural transmission and/or gene flow. Here we examine the robustness of song divergence to contact using cline analyses of molecular and phenotypic data.

## Study taxa

*Nectarinia* (née *Cinnyris*) *moreaui* and *N. fuelleborni* (alternately classified as the subspecies *N. mediocris fuelleborni*) are sunbirds of Eastern Afromontane forest and forest edges. Bowie et al. (2004) found that *moreaui* and *fuelleborni* possess 4-6% mtDNA_sequence divergence, and that both mtDNA lineages are found within forest habitats of the Udzungwa Mountains, Tanzania (see also [23]). The Udzungwa, much like the other Eastern Arc Mountain blocks (e.g. Ukaguru, Usambara, Uluguru), had largely been considered a single biogeographic ’island’ for forest-dependent birds [23,24,25], such that the existence of two highly divergent mtDNA lineages within the northeastern Udzungwa was not expected despite reports that individuals “within the concept” of *moreaui* had been captured there [26].

The forest areas inhabited by both species are constant environments that persisted in a similar state through the climatic fluctuations during the Plio-Pleistocene [27]. The build-up of avian paleo- and neo-endemic diversity in the northeast Udzungwa provides further evidence that these areas have maintained stable environments through time [24,25,28]. In contrast to contact zones where inter-specific interactions have been attributed to recent anthropogenic changes to local environments, the *moreaui-fuelleborni* contact zone occurs in tracts of mature forest in an area with high environmental stability.

## Methods

### Molecular methods – sampling, laboratory procedures, and molecular analyses

Tissue and blood samples were acquired from multiple field expeditions and museum collections. We sampled 132 individuals encompassing 14 different populations (Table S5). DNA extraction was performed using Qiagen DNEasy blood and tissue kits. Sanger sequencing was performed for the mtDNA gene ND2, the Z-linked intron loci CHDZ, MUSK, and BRM, and the autosomal introns 11836 and 18142 [29-33] (further details in Supplementary Materials). Sequences were checked for quality and aligned using the MAFFT alignment algorithm [34] within Geneious Pro 5.1.6 (2010). Nuclear introns were phased probabilistically into haplotypes using the program PHASE [35]. Population structure and individual population membership were investigated using the Bayesian population assignment algorithm of the program STRUCTURE [36]. The number of populations *k* was set *a priori* to 2 based on initial results where *k* was allowed to take values 1 to 8. Results of Bayesian population assignment are depicted from a) a structure run where only nuclear intron haplotypes were included, compared against mtDNA haplotypes scored as either *fuelleborni* or *moreaui* in origin; and b) a structure run where nuclear intron and mtDNA haplotypes were both included for each individual in the data set.

To provide an indication of the level of within-species population structure, multi-locus K_st_ indices [37] were calculated in DNAsp [38]. Individuals with mixed ancestry, as evidenced by low probability of assignment (<90%) to both *N. moreaui* and *N. fuelleborni* in structure analyses, were excluded from these analyses. Only populations with sampling of five or more individuals were included in this analysis, and only those from within the cline-fitting area were included, which resulted in Misuku being excluded.

### Song recording and analysis

Sound recordings were made in the field in 2008-10 using Sennheiser ME-67 shotgun microphones and Marantz solid-state recorders (models PMD660, PMD670, and PMD671; 16 bit precision, 44.1 or 48 kHz sampling rate). Two additional *N. fuelleborni* recordings were obtained from the Macaulay Library, Cornell University Lab of Ornithology. Values for song variables were obtained for each of 14 measurements from calculations based on data extracted for each unique element in Luscinia ([39], see Supplementary Materials for measurement definitions and procedures). Mean individual values were then calculated from the set of songs measured for each individual, since the individual is our unit of analysis. Each vector of mean individual values was scaled by its standard deviation in R [40] before principal components analysis in JMP 9 [41]. The first principal component was used in cline analyses.

Analyzed song number per individual varied from 1 – 30 songs (mean=3.3, SD=3.2), which were exclusively sampled from songs sung in bout form (i.e. multiple songs were sung consecutively with a fairly regular inter-song interval). Some individuals were banded prior to sound recording, and were identified by band combination. For unbanded individuals, a minimum distance of 60m between recording localities was used to establish separation between individual samples. This minimum distance threshold was selected based on extensive observation of territorial behaviour, and from observations of the typical densities of territorial males.

### Bill measurements

Culmen length, defined as the distance between the distal tip and the notch where the culmen meets the skull, was measured for male museum specimens using digital calipers. A single observer (JPM) measured 141 male bills. We retained the 122 measurements derived from populations along the cline transect (Figure 1) for cline analyses.

**Figure 1:**
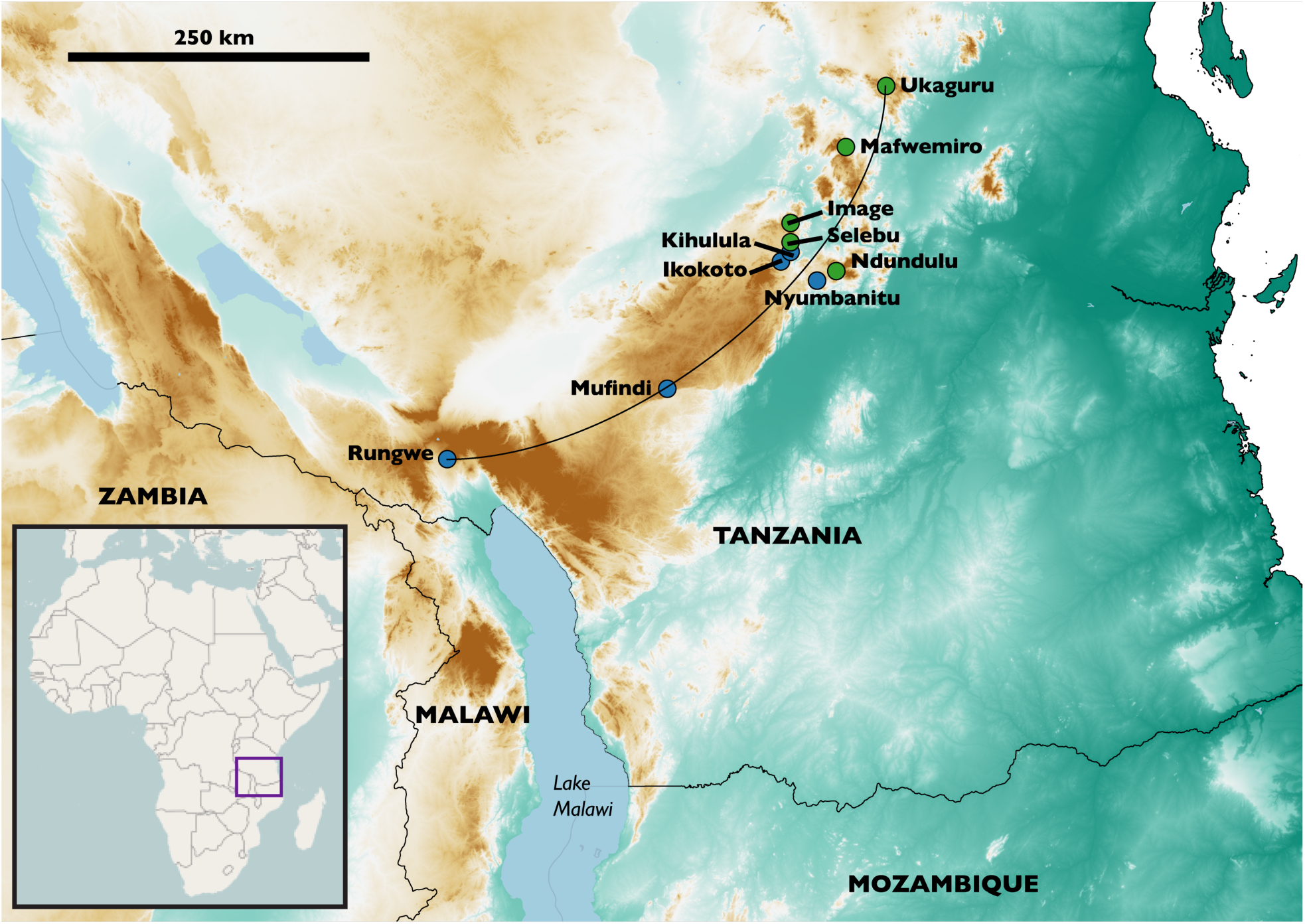
Map of the study area, with named localities mentioned in the text. The circular arc transect shown was used to reduce the spatial dimensionality of the data for cline analyses. Colouration indicates elevation: green shades = lower elevation and brown shades = higher elevation. Green dots are sampling localities dominated by *N. moreaui.* Blue dots are sampling localities dominated by *N. fuelleborni.*

### Cline analyses

To analyze individual- and population-level geographic variation in genotypes and phenotypes, we first reduced the spatial dimensionality of the data by assigning each population a position along a circular arc transect. We fixed the northernmost and southernmost sampling populations as the endpoints of the circular arc, and then adjusted the arc’s radius in ArcMap 10 [42] such that it evenly split the set of populations at the center of the contact zone (Figure 1). Population positions in one dimension were then assigned by finding the nearest point along the 504.5 km arc for each sampled population.

The goals of cline analyses were two-fold: to estimate the center and width of molecular and phenotypic clines as parameters of cline models, and to test for coincidence and concordance among molecular and phenotypic clines. Using the R package HZAR [43], we fit and compared clines for three traits: a molecular hybrid index (q-scores from STRUCTURE analyses of nuclear molecular variation), song PC1 scores, and culmen length (see Supplementary Materials for additional cline fitting details).

In HZAR, clines were fit with five different possible architectures: a sigmoid cline without exponential tails, the two models with a sigmoid cline and a single flanking exponential tail, a central sigmoid cline with identically parameterized exponential tails, and a central sigmoid cline with two independently fit exponential tails [11,43-46]. We then compared models representing each of the five architectures, using model selection by AICc to determine a single preferred model architecture for each trait. We then extracted the maximum likelihood model for each trait. For each architecture-trait combination, we made initial model fits (chain length = 1x10^6^ generations, burn-in = 5x10^5^ generations) to optimize the covariance matrix used in three separate subsequent MCMC chains [43]. These three secondary MCMC chains were then concatenated (9x10^6^ generations) for postprocessing and analysis. We checked for adequate mixing and convergence in cline parameter estimation by visualizing sampling trajectories with the R package coda [47].

### Niche similarity test

To test the hypothesis that *moreaui* and *fuelleborni* have diverged under conserved abiotic niches, we performed a one-tailed background similarity test of modeled bioclimatic niche [48]. The background similarity method tests the hypothesis that projected ecological niche models developed from the spatial distribution of two biological entities exhibit a level of similarity that differs from the similarity of projected niches modeled from randomly sampled points within a buffer area around occurrence points. This buffer area should include areas that would likely be encountered by the focal organisms via dispersal over relevant evolutionary timescales. To generate the null distribution, a series of psuedo-randomly sampled points from the buffer area (“background samples”) surrounding species A’s occurrences are used to generate a niche model that is compared to the niche model generated for the actual occurrence points for species B. For this test, we used a buffer zone of 100 km around the set of distribution points for each species (see Supplementary Materials).

Niche models from both the true occurrences and 100 sets of background sample points from the *N. fuelleborni* and *N. moreaui* buffers were generated using the maximum entropy distribution modeling software Maxent and a set of seven BIOCLIM variables [49-51]. Maxent uses presence data and background samples (distinct from points sampled from the buffer area for the background similarity test) to produce predictive niche models by machine-learning maximum entropy modeling. The generated model is the maximum entropy distribution constrained only by the relationships of the features (the BIOCLIM variables) in predicting the occurrence data against the environmental background [51].

## Results

### Molecular sequence data

Of the six molecular loci, *moreaui* exhibited a statistically significant K_st_ (with Bonferroni correction) among sampled populations only for MUSK (Table S1). The minimal structure in *moreaui* was unexpected given that sampled populations are isolated by unsuitable low-elevation environments, and that many bird species with similar distributions exhibit strong structure among the same mountain blocks [23,24,52]. This result suggests that dispersal has been important in the population history of *N. moreaui. N. fuelleborni* exhibited statistically significant K_st_ among populations for ND2 and MUSK. Population structure for these loci was driven by differences between the Rungwe and Udzungwa (Mufindi, Ikokoto, and Nyumbanitu) populations, which are separated by the Makambako Gap, a known biogeographic break [53].

Population assignment in STRUCTURE closely matched *a priori* expectations for species identity of individuals based on morphology, geography, and previous mtDNA phylogeography [12]. We use a cutoff value of .90 posterior probability of assignment to designate ’pure’ individuals of either species from STRUCTURE output. Individuals were dichotomously scored for mtDNA species origin by inspecting the alignment (Figure 2). In the combined mitochondrial and nuclear sequence data set, seven of 132 individuals show mixed ancestry. Five individuals have recombinant nuclear genomes, and two individuals exhibit cyto-nuclear discordance. The seven individuals of mixed ancestry come from four populations: Kihulula, Nyumbanitu, Image, and Ndundulu (Figure 1). Sympatry of pure parentals species is revealed from molecular analyses at only two localities: Nyumbanitu and Ikokoto (Figures 1 and 2).

**Figure 2:**
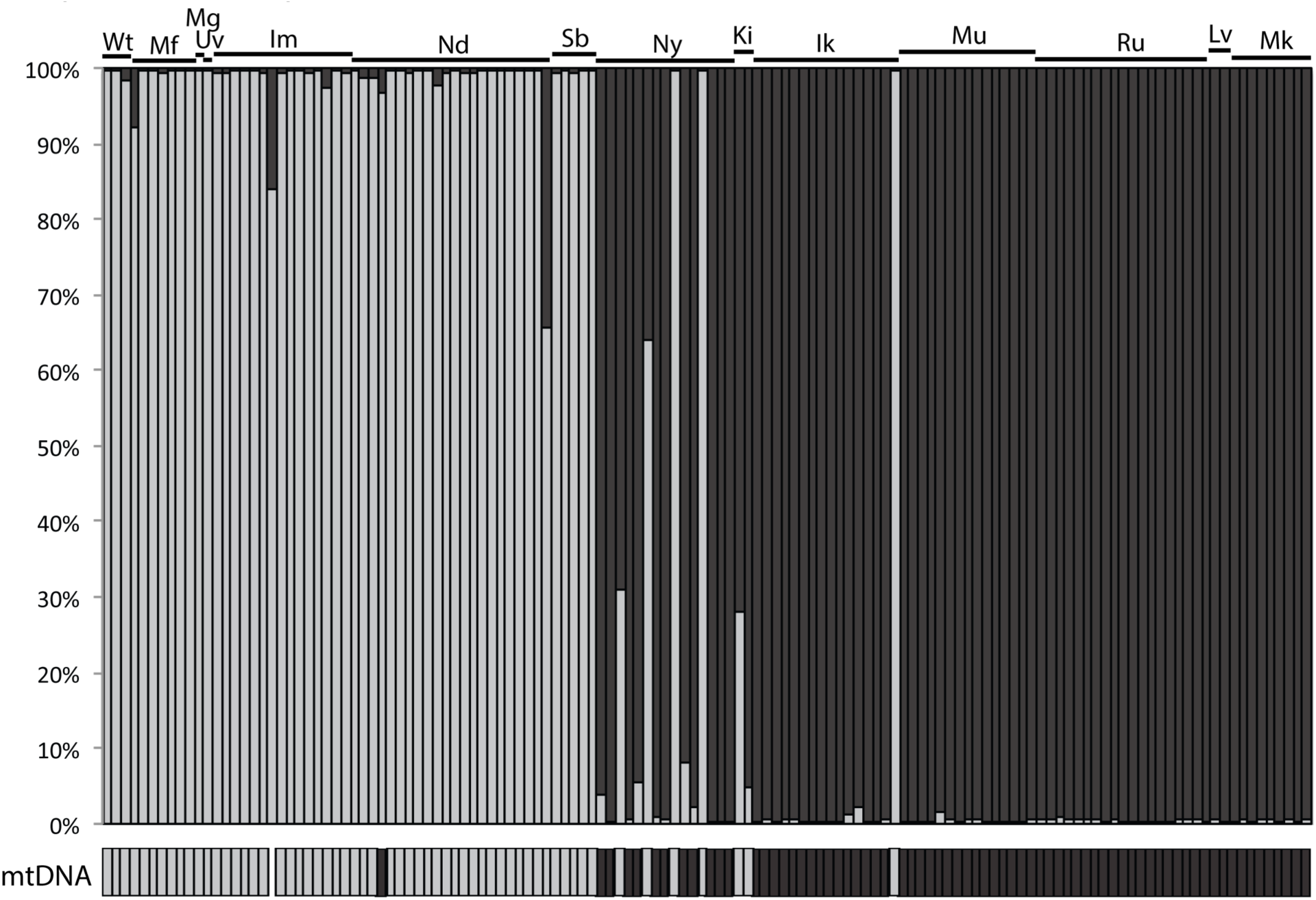
Probability of assignment to *N. moreaui* (light gray) and *N. fuelleborni* (dark gray) as determined from a STRUCTURE analysis using nuclear gene haplotypes (top). The corresponding mtDNA lineage (*moreaui* or *fuelleborni*) for each individual is also shown (bottom). Note two instances of mito-nuclear discordance. Population abbreviations are as follows: Wt = Wota, Mf = Mafwemiro, Mg = Mang’alisa, Uv = Uvidunda, Im = Image, Nd = Ndundulu, Sb = Selebu, Ny = Nyumbanitu, Ki = Kihulula, Ik = Ikokoto, Mu = Mufindi, Ru = Rungwe, Lv = Livingstone Mountains, Mk = Misuku Hills.

### Species-level phenotypic divergence

*Songs* Representative sonograms of the two species can be found in Fig. 1 of McEntee (2014). MANOVA following scaling and centering of all 14 variables was significant for differences between allopatric populations of the two species (n_1_ = 38 *fuelleborni, n_2_* = 41 *moreaui,* F = 268.87, df = 1, *p* < 1 x 10^−15^). The first principal component corresponded strongly with species differences. Loadings for PC1 for all nine variables with statistically significant differences in the individual ANOVAs (Tables S2 and S3) had absolute value >.45, indicating correlation among these variables along the axis of species differences.

*Bills* Mean ± SE male bill lengths were 23.29 ± 0.15 for *N. fuelleborni* and 24.89 ± 0.11 for *N. moreaui* (*n*_1_ = 51 *fuelleborni, n_2_* = 92 *moreaui,* t = -8.86, df = 97, *p* < 1 x 10^−13^; individuals with mixed ancestry were excluded from this comparison.

*Ecological niche* Species distribution models from Maxent had high prediction value for the training set for both species (*fuelleborni* AUC: .966; *moreaui* AUC: .988). In relativized contribution and permutation tests performed using Maxent, temperature variables had a higher impact than precipitation variables. Predictions projected into the range of the sister species resulted in remarkably high concordance between the projected niche and the sister species’ localities (Figure 3a).

**Figure 3:**
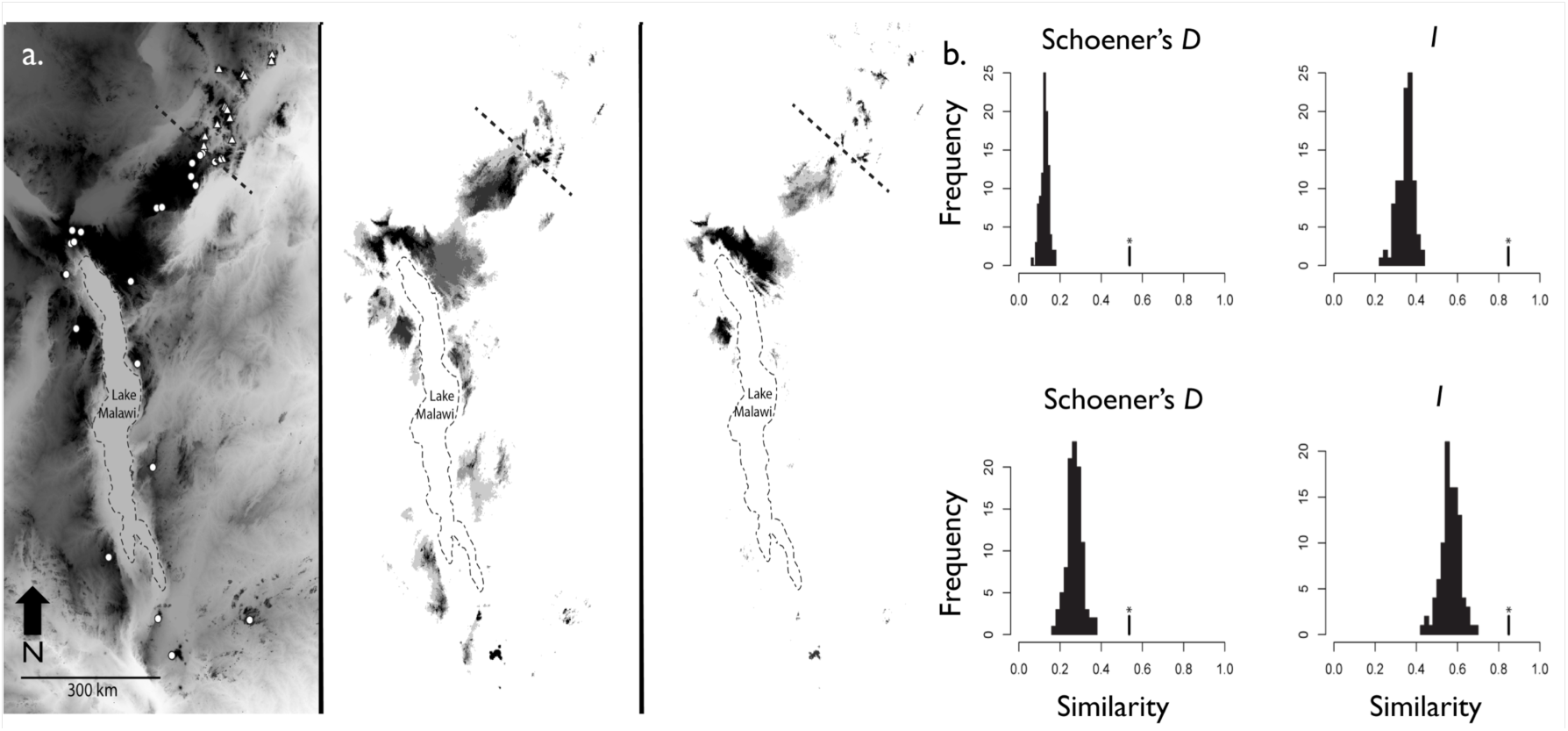
a) From left to right: occurrence points for *Nectarinia moreaui* (triangles) and *N. fuelleborni* (circles) with elevation shown across their distributions (darker colours at higher elevations); Maxent niche model for *fuelleborni,* with darker colour indicating higher suitability; equivalent model for *moreaui.* b) Results of background similarity tests performed in ENMTools. Starred bars for the niche similarity indices Schoener’s *D* and *I* are similarity scores from niche models predicting the distribution of the other species (top: *fuelleborni* model predicting *moreaui* suitability, bottom: *moreaui* model predicting *fuelleborni* suitability). Distributions shown are null distributions generated from niche models developed for sets of ‘background’ points within 100 km buffers around occurrence points. The greater similarity seen in niche models generated from the actual sister taxa distributions as compared to the null distributions indicates strong niche conservatism. See Methods for additional details.

Results of the background similarity test, performed in ENMTools [54], are shown in Figure 3b. Compared to the null distributions of niche similarity scores generated from comparing background point models of species A with the actual distribution model for species B, niche models for *fuelleborni* and *moreaui* had higher similarity scores for both ecological indices calculated: Schoener’s *D* and *I.* These tests indicate abiotic niche conservatism through speciation, with niche similarity greater relative to background similarity (*p* = .01) for all four comparisons.

## Clines

The preferred cline architecture for culmen length included no flanking exponential tails, while the preferred cline architectures for the molecular index and song clines both included exponential tails (Table 1). The model including only a right tail was the preferred model for the molecular index. Among song cline models, the left-tail only, right-tail only and two-identical-tail models had similar AIC scores (∆AICs < 0.32). Cline center and width estimates for the molecular index and song clines were coincident and concordant, with center ~163 km and width ~6 km. The center of the molecular and song clines corresponds closely to the positions of Nyumbanitu and Kihulula forests along the transect.

**Table 1.** Center and width estimates with 95% confidence intervals for each of the five modeled cline architectures used to investigate each of the three examined traits. For each trait, preferred models (ΔAIC<2) are in bold.

| Trait | Center | Width | $\Delta AIC$ | Tails | Parameters |
| --- | --- | --- | --- | --- | --- |
| Song PC1 |  |  | 3.88 | none | 7 |
| **Song PC1** | **163.0 (161.3-165.0)** | **5.9 (0.8-9.5)** | **0** | **left** | **9** |
| **Song PC1** |  |  | **0.31** | **right** | **9** |
| **Song PC1** |  |  | **0.14** | **mirror** | **9** |
| Song PC1 |  |  | 5.07 | both | 11 |
| Molecular index |  |  | 15.95 | none | 7 |
| Molecular index |  |  | 17.30 | left | 9 |
| **Molecular index** | **163.7 (159.1-164.8)** | **3.7 (0.3-5.5)** | **0** | **right** | **9** |
| Molecular index |  |  | 11.57 | mirror | 9 |
| Molecular index |  |  | 5.21 | both | 11 |
| **Culmen length** | **155.1 (140.4-158.8)** | **21.4 (8.6-54.0)** | **0** | **none** | **7** |
| Culmen length |  |  | 4.54 | left | 9 |
| Culmen length |  |  | 5.24 | right | 9 |
| Culmen length |  |  | 9.78 | mirror | 9 |
| Culmen length |  |  | 9.80 | both | 11 |

The culmen length cline is not coincident with the molecular and song clines. The 95% credible interval for the maximum likelihood culmen length cline center does not overlap with the 95% credible intervals for the maximum likelihood molecular index and song centers. The culmen length cline’s center is shifted ~8 km north of the molecular index and song cline centers. The culmen length cline is also broader in width than the molecular index cline (95% credible intervals do not overlap), while there is a small amount of overlap between the 95% credible intervals for culmen length and song cline widths.

## Discussion

### Evidence for social selection parapatry

Models indicate that (incipient) species with similar, even identical, ecological niches may establish stable mutual species borders, and may thus persist for long time periods within a finite domain [55]. West-Eberhard’s [6] “social selection parapatry” hypothesis is an appealingly plausible explanation for how such stable borders could arise. Demonstrating “social selection parapatry” minimally requires satisfying the following criteria: 1) demonstrating that each species’ range is not predominantly limited by aspects of the environment outside of the presence of the sibling species, for example by a geographic barrier to dispersal or an abiotic gradient [3]; 2) showing that ecological competition is intense where there is local co-occurrence; 3) demonstrating that selection against hybrids is insufficient to account for the totality of fitness loss for each species at the parapatric edge of its range, and 4) establishing that social selection is the primary mechanism for phenotypic divergence. We review the evidence for each of these components from the *moreaui* - *fuelleborni* parapatric boundary.

Neither a geographic barrier nor an abiotic gradient can explain the range border of either species in the northeast Udzungwa. Though montane forests occurring at 1400 m and higher, where the two species breed, are patchily distributed on the landscape in the vicinity of the range boundary, the short distance to suitable habitat at each species’ range edge precludes the possibility that a geographic barrier is responsible for limiting the range of either species. Moreover, there is local syntopy and hybridization, thus the two species are clearly not fully separated in space. An abiotic gradient, meanwhile, is extremely unlikely to be more important than the presence of the heterospecific in limiting each species’ range. The results of the niche models and background similarity tests presented here (Figure 3) indicate that the environment is suitable for each species well beyond the range edge, extending far into the range of the sibling species. We acknowledge that we have not tested an additional possibility, that biotic factors outside of the sibling species interactions (e.g. plant food availability) play a major role in limiting the range of either species. However, the striking ecological similarity of the two species suggests that their populations are unlikely to be independently regulated. With respect to food plant species, both species are generalists in their exploitation of nectar ([56], JPM pers. obs.), and both frequently feed on the same widespread plants (*Tecoma capensis*, *Halleria lucida*, *Dombeya* sp., *Leonotis spp.,* JPM pers. obs.) in the vicinity of the parapatric boundary. Consequently, we conclude that the range of each species is limited primarily by the presence of the sibling species, and not by aspects of the abiotic environment.

For the second criterion, that the two species are in strong ecological competition where they locally co-occur, our evidence comes from a previous simulated territorial intrusion study [57] and from natural history observation. In the simulated territorial intrusion study, populations of both species near the species border exhibited strong aggressive responses to heterospecific mounts following approaches initiated by heterospecific or conspecific song playback [see Figure 5 in 57]. This result suggests that interspecific interference competition for territories should be strong where there is local occurrence. At the only site where pure individuals of both species hold adjacent territories, we have observed extended aggressive chases among males of the two species. During these chases, males used vocalizations that are known otherwise only from aggressive intra-specific interactions (pers. obs., JPM, MM, EM), suggesting that heterospecifics can elicit similar aggression as conspecifics, and therefore that inter- and intraspecific interference competition may be of similar magnitude. Territorial disputes such as this are further suggestive of resource competition, where suitable breeding territory is a shared resource. Collectively, the experimental and observational evidence is consistent with the social selection parapatry hypothesis, which requires strong ecological competition between species.

With respect to the third criterion above, the combination of incomplete reproductive isolation and selection against hybrids can generate Allee effects for the locally rare species via reproductive interference [58] near mutual species borders [3]. It is theoretically possible then for reproductive interference to limit species ranges even where ecological competition is not strong. However, because reproductive interference likely combines with ecological competition to set Allee effects in nature, teasing out their relative effects is likely to be difficult. The strongest support for West-Eberhard’s social selection parapatry hypothesis, which specifies ecological competition and less so reproductive interference as the driver of mutual exclusion, would come from parapatric distributions where no reproductive interference occurred. In this study, individuals of mixed ancestry comprise 7 of the 75 individuals (9.3%) sampled from sites where genes from both species were sampled (Figure 2). While this proportion is likely to be an overestimate of contact zone hybrid formation because of intensive sampling effort where syntopy of parental forms was discovered, it is nonetheless clear that pre-zygotic reproductive isolation is incomplete. We must then conclude that reproductive interference via selection against hybrids is likely a contributor to the range limits of *fuelleborni* and *moreaui* at their shared range boundary. However, because evidence indicates strong ecological similarity between species, reproductive interference and ecological competition can be seen as combining to limit the range of each species. Teasing out the relative strength of effects of reproductive interference and ecological competition on range limits would be a productive avenue of future research [59,60]. Of the four examined criteria for social selection parapatry, the insufficiency of reproductive interference to explain range limits remains the most challenging to support with evidence. It is, moreover, surprising that birds with such different songs do not exhibit more complete reproductive isolation. Even dramatic divergence in social traits is insufficient then to prevent hybridization attempts, such that reproductive interference could play a strong role in stabilizing avian parapatric boundaries even where (attempted) hybridization has not yet been documented.

With respect to the last criterion above, we show that a social trait, male territorial song, is highly divergent between *moreaui* and *fuelleborni,* whereas other traits (bill morphology, bioclimatic ecological niche) are more subtly divergent. Here we assess the evidence that each of the following mechanisms has contributed to the described song divergence: acoustic adaptation, byproduct mechanism from natural selection on other traits (e.g. morphology that constrains sound production; [61]), genetic and/or cultural drift [17,62], and social selection independent of environmental differences experienced by populations. We argue that, of these possibilities, social selection independent of ecological or morphological differences is the most plausible primary mechanism for song divergence between *moreaui* and *fuelleborni.* However, the action of social selection together with genetic drift may be important, as suggested by models incorporating both processes [63,64].

The “acoustic adaptation hypothesis” applied to species-level divergence requires that there is little overlap of the acoustic window experienced by divergent species. The background similarity tests presented here provide evidence that the focal species use extremely similar bioclimatic niche space, while natural history observations indicate the two species sing from and otherwise make use of a broad variety of forest and forest edge microhabitats, suggesting extensive overlap in the acoustic window each species experiences. The avian community context, another source of acoustic window constraint variation, varies across space. A single species, *Serinus whytii,* occurs throughout much of *fuelleborni*s range but is absent from *moreaui* s, and its vocalizations could impinge on the acoustic window of *fuelleborni.* However, instead of showing divergence from *S. whytii* as expected from_limiting of the acoustic window, *fuelleborni* sings songs that are similar to *S. whytii*’s songs in gross structure (JPM, unpublished data, see also [57]). Overall, the constraints of the acoustic window are likely to be quite similar for both *moreaui* and *fuelleborni.* Mediation of divergence by shifted acoustic constraints is then unlikely to contribute substantively to song divergence.

The “byproduct hypothesis” requires phenotypic change in a trait with pleiotropic effects on song. Multiple studies have shown that bill morphology divergence is associated with song divergence [65,66]. The mean of *moreaui*’s culmen lengths is greater than that of *fuelleborni,* providing evidence that pleiotropic effects on trills are possible. However, one prediction of the byproduct hypothesis for song is that “ecologically” selected traits and song must co-vary among populations. Cline analyses show that *moreaui* populations at Ndundulu and Selebu sing species-typical songs despite possessing smaller (more *fuelleborni*-like) bills relative to other *moreaui* populations, indicative of a subtle decoupling of bill length and song phenotype variation that runs counter to the prediction of the byproduct hypothesis that song and bill phenotypes should tightly co-vary among populations ([18], Table 2). The dramatic divergence in song duration (Table S2) and fine-scale spectro-temporal aspects of song structure (e.g. coefficient of variation in interval duration, Table S2) cannot easily be explained by pleiotropic effects of bill divergence. Our study thus discounts a strong role for bill morphology in song divergence between *moreaui* and *fuelleborni*.

Random genetic drift can explain song divergence if different alleles underlying song variation fix in different species from ancestral standing genetic variation. A prediction from this genetic drift model is isolation-by-distance in song phenotype aspects that are vertically inherited. While we do not specifically test this hypothesis here, we note that the song phenotype dimension presented (PC1) is strongly conserved across space within species (Figure 4) while being strongly divergent among species. This pattern suggests that different processes are at play within versus between species [67], which is confirmed in a separate study specifically examining population-level song variation (McEntee et al. in prep). Strong conservatism in song phenotypes among spatially isolated populations, including some that have accrued substantial divergence at molecular markers (Table S1, McEntee et al. in prep), suggests that stabilizing selection prevents random genetic drift alone from driving song phenotype divergence among population isolates within species. Cultural drift, meanwhile, could explain the use of different song elements among populations within species, but cannot account for the gross structural divergence found in *moreaui* and *fuelleborni* song (Table S2).

**Figure 4:**
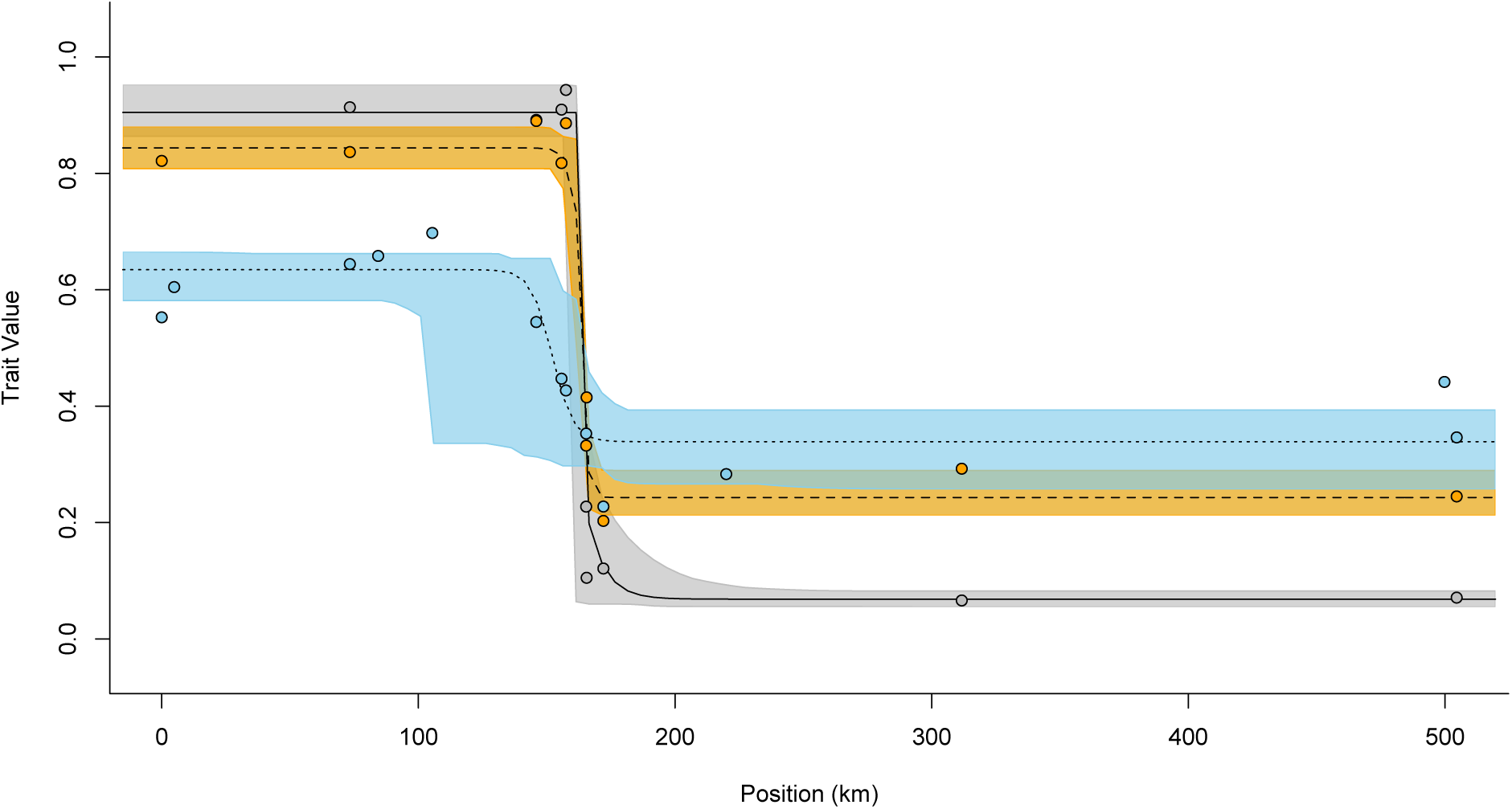
Cline analysis (HZAR) results for the molecular hybrid index (gray), song PC1 (orange), and culmen length (blue). Lines indicate the maximum likelihood cline for each trait, from the model with the minimum AICc value for that trait. Shaded areas represent 95% confidence intervals. Points are mean values for populations included in the cline analyses. When the clines are restricted to a common scale, the culmen length cline occupies the narrowest range of trait values because the maximum and minimum individual culmen length values lie far from the population means.

The failure of the acoustic adaptation, byproduct, and drift mechanisms to explain song divergence in this case leaves social selection independent of environmental differences as the most plausible primary mechanism of divergence. What is perhaps most interesting about song divergence in this case is within-species conservatism despite dramatic divergence between *moreaui* and *fuelleborni* (Figure 4). Conservatism is suggestive of stabilizing selection [68,69]. However, if stabilizing selection on song prevents among-population divergence within species, what processes can explain rapid divergence between species? We suggest range expansion-associated divergence as a possibility [70]. We hypothesize that *N. fuelleborni* arose as a daughter species of *N. moreaui* during an ancient southward range expansion, with an attendant abrupt shift in song phenotype, and that the two have subsequently come into secondary contact, or perhaps have persisted in contact for a long period of time. This hypothesis could account for both the striking divergence in song between *fuelleborni* and *moreaui* and the comparatively limited geographic variation within each species.

### Robustness of learned song divergence to secondary contact and hybridization

The role of learned song in bird speciation remains an open question that falls into the larger debate regarding plasticity’s importance in divergence [71,72]. The combination of learned song’s lability and its role in mate choice suggests it could promote speciation by reducing or impeding gene flow. However, the song learning process can result in heterospecific copying [73,74]. Divergent learned song may then be a weak barrier to hybridization [19,20,75-77]. There is more potential for learned song differences to prevent gene flow if song pre-dispositions and adult song phenotypes have co-diverged with preferences or response functions. Experiments have shown that certain aspects of song phenotypes tend to be constrained by innate song predispositions, namely duration, frequency range, organization (gross structure), and element (syllable) morphology [78-82]. Thus (incipient) species bearing divergence in these aspects are less likely to converge on contact. *N. moreaui* and *fuelleborni* song differences include divergence in all of the listed aspects likely to be constrained by innate predispositions, rendering these species’ contact zone an important case study for how song predisposition divergence impacts the robustness of learned song divergence to secondary contact with hybridization.

The coincidence and concordance of molecular and song clines (Tables 1 and S4, Figure 4) suggest that song behaves much as a quantitative trait under within-species stabilizing selection across the *N. moreaui* – *N. fuelleborni* contact zone, and indeed suggests that heterospecific copying has not broadened the song cline, as would be predicted from inter-specific cultural transmission in parapatry. It is important to note, though, that not all intermediate values for learned song come from mixed-ancestry individuals. At least one of the intermediate values for song in the contact zone was exhibited by a pure *moreaui* individual. Therefore, those individuals with intermediate song phenotypes may represent hybrids as well as pure individuals. Thus, in the geographic context analyzed here, song behaves much like a typical quantitative trait, but heterospecific copying is apparent within one contact zone population. If heterospecific copying is disadvantageous [75], selection against locally rare heterospecifics exhibiting mis-matched genotypes and phenotypes might complement selection against hybrids in reinforcing the stability of the cline.

## Conclusions

We have shown that *N. moreaui* and *fuelleborni* may be viewed as a case of social selection parapatry. Our evidence includes demonstrating 1) that the two species have diverged strongly in a social trait (learned song) while showing strong conservatism in bioclimatic niche, 2) that interspecific interactions, not an ecological gradient or geographic barrier, play a primary role in limiting the range of each species, and 3) that ecological competition is intense where there is syntopy. While interspecific competition is apparent and clearly plays some role in limiting the range of each species, we cannot yet rule out the possibility that reproductive interference in the form of selection against hybrids (and potentially mate-searching inefficiency for the locally rare species) is essential in restricting the range of each species. The relative contribution of ecological competition and reproductive interference in limiting the ranges of parapatric species may be difficult to distinguish, but doing so should be a focus of future research.

The abrupt spatial turnover in highly divergent song phenotypes, concordant with genotypes, recommends the *moreaui*-*fuelleborni* hybrid zone for future study on song divergence and speciation. Strong divergence in multiple aspects of song phenotype in these two species, together with the possibility of examining the phenotypes of hybrids, may allow for identification of the molecular changes underpinning divergent song predispositions.

## Acknowledgements

We owe thanks to Norbert Cordeiro,_Liz Baker and Neil Baker for advice on field work. We thank the late Jacob Kiure for his field sampling efforts in Tanzania. Gary Voelker, Thomas Gnoske and Potiphar Kaliba are thanked for assistance in the field in Malawi and the Field Museum of Natural History is thanked for loan of tissues. For assistance with the cataloguing of sound recordings, we thank Nadje Najar, Emilia Wakamatsu, Addien Wray, Jessica Hughes, Somin Lim, Violet Kimzey, and Kyle Marsh. David Taylor, Jenny Woodier, Sarah McGregor, and Shanna Sheridan-Johnson provided lodging during field work. Permissions for Tanzanian field research were furnished by COSTECH (Commission for Science and Technology), TAWIRI (Tanzania Wildlife Research Institute), and the Division of Forestry and Bee-keeping. JPM thanks Craig Moritz, Frederic Theunissen, and the “Endler reading group” for critical discussion and advice in the development of the study. Robert Lachlan advised regarding the use of his sound analysis program Luscinia. Kimball Garrett assisted with access to museum specimens at the Los Angeles County Museum. Sandra Perello assisted in the measuring of museum specimens. Lydia Smith and Anna Sellas contributed to the molecular aspects of the study._

## Data accessibility

Molecular data will be contributed to GenBank. Song and bill measurements will be made accessible via the Dryad Digital Repository datadryad.org. Song recordings will be made available through the ARCTOS data portal.

## Ethics

Specimens and blood samples were acquired in accordance with the laws of Tanzania and Malawi following ethical guidelines suggested by the American Ornithologists Union. Protocols were reviewed and approved by a University of California, Berkeley, IACUC.

## Authors’ Contributions

JPM designed the study, conducted field research, performed molecular analyses, analyzed data, and wrote the manuscript. JVP performed molecular analyses. CW facilitated and conducted field research. EM and MM conducted field research. LH and JF organized and conducted early field research, laying the foundation for this study, and edited the manuscript. RCKB designed the study, conducted field research, analyzed molecular data, and wrote the manuscript. All authors gave final approval for publication.

